# DomBpred: protein domain boundary predictor using inter-residue distance and domain-residue level clustering

**DOI:** 10.1101/2021.11.19.469204

**Authors:** Zhongze Yu, Chunxiang Peng, Jun Liu, Biao Zhang, Xiaogen Zhou, Guijun Zhang

## Abstract

Domain boundary prediction is one of the most important problems in the study of protein structure and function, especially for large proteins. At present, most domain boundary prediction methods have low accuracy and limitations in dealing with multi-domain proteins. In this study, we develop a sequence-based protein domain boundary predictor, named DomBpred. In DomBpred, the input sequence is firstly classified as either a single-domain protein or a multi-domain protein through a designed effective sequence metric based on a constructed single-domain sequence library. For the multi-domain protein, a domain-residue level clustering algorithm inspired by Ising model is proposed to cluster the spatially close residues according inter-residue distance. The unclassified residues and the residues at the edge of the cluster are then tuned by the secondary structure to form potential cut points. Finally, a domain boundary scoring function is proposed to recursively evaluate the potential cut points to generate the domain boundary. DomBpred is tested on a large-scale test set of FUpred comprising 2549 proteins. Experimental results show that DomBpred better performs than the state-of-the-art methods in classifying whether protein sequences are composed by single or multiple domains, and the Matthew’s correlation coefficient is 0.882. Moreover, on 849 multi-domain proteins, the domain boundary distance and normalised domain overlap scores of DomBpred are 0.523 and 0.824, respectively, which are 5.0% and 4.2% higher than those of the best comparison method, respectively. Comparison with other methods on the given test set shows that DomBpred outperforms most state-of-the-art sequence-based methods and even achieves better results than the top-level template-based method.

## 1 Introduction

Protein structure domains are the fundamental units of protein structure, folding, function, evolution and design. More than 80% of eukaryotic proteins and 67% of prokaryotic proteins contain multiple domains, and many biological functions rely on the interactions amongst different domains [1, 2]. Thus, deducing protein domain boundaries is an essential step for determining structural folds, understanding biological functions and/or annotating evolutionary mechanisms [3]. Furthermore, with the revolutionary success of AlphaFold2 in protein structure prediction [4], there is a consensus in the community that the protein structures prediction for single domain is nearly solved [5]. In CASP14, AlphaFold2 structures have a median backbone accuracy of 0.96 Å RMSD95 (C_α_ root-mean-square deviation at 95% residue coverage, 95% confidence interval = 0.85–1.16 Å) [4], where CASP assesses the quality of protein structure predictions primarily on the individual domains. However, the full-chain modelling of multi-domain proteins remains a challenge in the absence of co-evolution information. There are two main approaches in the current modelling multi-domain protein structures [6]. The first is to directly predict the full-chain model from the protein sequence, such as AlphaFold [7], RaptorX [8] and I-TASSER [9]. The second is to divide multi-domain proteins into single domains, and then assemble the separately modelled domain structures into full-chain models through domain assembly methods, such as DEMO [2] and AIDA [10]. As the length of the protein sequence increases, directly predicting the full-chain model becomes very difficult and inefficient, but the assembly method is almost unaffected by the protein full-chain length. Therefore, how to accurately predict the domain boundary is the basis and key of the domain assembly methods.

In general, domain boundary prediction methods can be classified into structure-based or sequence-based. Structure-based methods require experimental or predicted protein structures for domain identification. CATHEDRAL [11] compares target protein structure against the template structures derived from the CATH [12] database to detect domains. DomainParser [13] uses a graph-theoretic approach to predict the domain boundary, in which each residue of a protein is represented as a node of the network and each residue-residue contact is represented as an edge with a particular capacity depending on the type of the contact. PDP [14] and DDOMAIN [15] split proteins into domains depending on the assumption that there are more intra-domain residue contacts than inter-domain contacts. DHcL [16] decomposes protein domains by calculating a van der Waals model of a protein. SWORD [17] assigns structural domains through the hierarchical merging of protein units, which are evolutionarily preserved substructures that describe protein architecture at an intermediate level, between domain and secondary structure. Furthermore, some methods are based on 3D structure prediction. For example, RosettaDom [18] uses the Rosetta *de novo* structure prediction method to build three-dimensional models, and then applies Taylor s structure based domain assignment method to parse the models into domains. These methods which deduce domains from protein structures generally have higher accuracies, but they can be applied only to proteins with short sequence lengths or with known experimental structures due to the limited structure prediction ability and experimental limitations [3]. Compared with structural information, protein sequence information is easier to obtain [6]. Therefore, methods that predict domain boundary from sequences are more meaningful and challenging in principle [3].

Sequence-based domain identification methods can typically be categorised into two general groups. The first group is primarily homology-based methods, which detect domains by comparing them with homologous sequences having known annotated domains. For example, Pfam [19], CHOP [20] and FIEFDOM [21], in which the target sequences are searched through known protein structure or family libraries through hidden Markov model (HMM) or PSI-BLAST programs. Then, the domain boundary information is obtained from the homologous template or family. ThreaDom adopts a threading-based algorithm to improve remote homologous templates detection [22]. It firstly uses the LOMETS [23] program to thread a target sequence through PDB to find homologous templates followed by the multiple sequence alignment (MSA) construction based on the target sequence. According to these MSAs, a domain conservation score is calculated to measure the conservation level of each residue and further used to judge boundary regions. An extended version, ThreaDomEx[24], is further developed to assign discontinuous domains by domain-segment assembly. These homology-based methods can reach a high accuracy of predictions when close templates are identified, but the accuracy sharply decreases when the sequence identity between the target and template is low [22]. Another group of methods is *ab initio* methods, which can overcome this limitation to some extent [6], with representative examples including DOMPro [25], DoBo [26], ConDo [27], DNN-dom [28], and FUpred [3]. DOMPro [25] trains recursive neural networks for domain models with training features including sequence profiles, predicted secondary structure, and solvent accessibility. DoBo [26] introduces the evolutionary domain information that is included in homologous proteins into the protein domain boundary prediction. ConDo [27] utilises neural networks trained on long-range, coevolutionary features, in addition to conventional local window features, to detect domains. DNN-dom [28] combines a convolutional neural network and bidirectional gate recurrent unit models to predict the domain boundary of a protein. FUpred [3] predicts protein domain boundary utilising contact maps created by deep residual neural networks coupled with coevolutionary precision matrices. However, the accuracy of sequence-based *ab initio* methods is often not high, although they are more practical especially in the absence of homologous sequences [3].

In the present work, we propose a novel *ab initio* sequence-based method, named DomBpred, to predict the domain boundary from protein sequences. In DomBpred, we construct a comprehensive domain sequence database based on the SCOPe and CATH databases and design an effective sequence metric to classify single-domain protein and multi-domain protein. For the multi-domain protein, a domain-residue level clustering algorithm inspired by Ising model is proposed to cluster the spatially close residues according to the inter-residue distance. The unclassified residues and the residues at the edge of the cluster are tuned by the secondary structure to form the potential cut points. Finally, a domain boundary scoring function is proposed to recursively evaluate the potential cut points to determine the domain boundary. Experimental results show that the domain classification ability and domain boundary prediction ability of DomBpred are better than those of comparison methods.

## 2 Materials and methods

The pipeline of DomBpred is shown in Figure 1. Starting from the input sequence, Jackhmmer [29] is employed to search homologous from a constructed single-domain sequence library (SDSL) to generate MSA, and then an effective sequence metric is used to determine whether the target sequence is a single-domain protein. If the target sequence is a multi-domain protein, the potential cut points are obtained by clustering close residues on the distance map. Then, the potential cut points are further tuned according to the predicted secondary structure information. Finally, a domain boundary scoring function is used to detect the domain boundary locations from the potential cut point collection.

**Figure 1.**
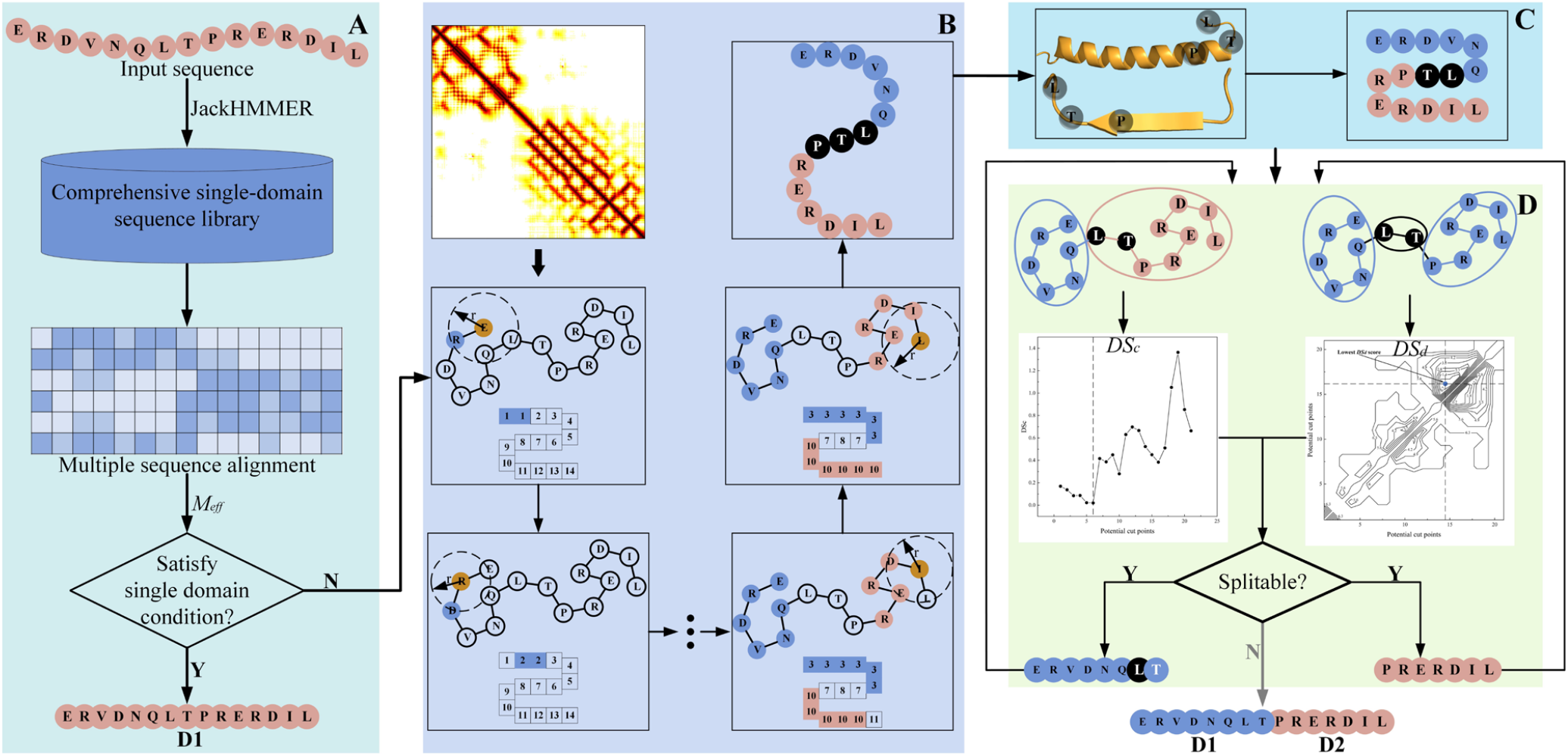
Pipeline of DomBpred. (A) Single- and multi-domain protein classification, where a sequence metric is used to detect the classification of the target sequence. (B) Domain-residue level clustering for potential cut points. A set of potential cut points is obtained by clustering spatially close residues in the distance map. (C) Cut points adjustment. The potential cut points are tuned based on the predicted secondary structure. (D) Domain boundary determination. The domain boundary scoring function is used to evaluate potential cut points, and the domain boundary is finally generated based on the cut points.

### 2.1 Single- and multi-domain proteins classification

Known domain knowledge can be used to assist the identification of single-domain and multi-domain proteins from sequence. Thus, we collect the sequences of individual domain from SCOPe 2.07 database [30] and CATH-Plus 4.3 database [31] to build a SDSL. In the data collection of SDSL, the protein sequence with more domains is included in SDSL if the same protein has a different number of domains defined in the two databases. If the same protein has same the number of domains defined in the two databases, the protein sequence defined in CATH-Plus 4.3 database is saved in SDSL. Finally, a pair-wise sequence identity < 100% and a sequence length > 35 residues are used to filter the domain sequences to obtain the final SDSL.

Based on the SDSL, Jackhmmer [29] program is used to generate a MSA for the target sequence. In module A of pipeline, an effective sequence metric is used to determine whether the input sequence is a single-domain protein. The effective sequence metric is defined as follows:

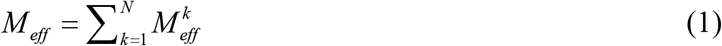

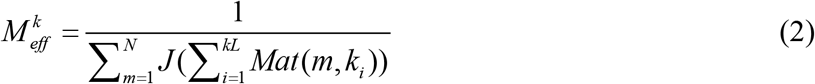

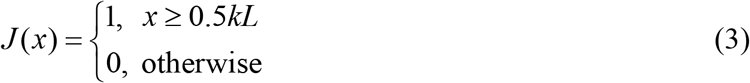

where *N* is the number of sequences in MSA and *kL* is the length of the *k*-th sequence in MSA; and *Mat*(*m, k*_*i*_) represents whether the *i*-th residue of the *k*-th sequence in MSA forms a matched residue pair with the residue of the *m-*th sequence. *Mat*(*m, k*_*i*_) = 1 if there is a matched residue pair; otherwise, *Mat*(*m, k*_*i*_) = 0. If the number of matched residue pair exceeds half of the length of the *k*-th sequence, the *m*-th sequence is considered as the similar sequence to the *k*-th sequence, and then *J* (*x*) = 1; otherwise, *J* (*x*) = 0. An illustration is shown in Figure 2. When *M*_*eff*_ is greater than 1, the input sequence is judged as a multi-domain sequence, otherwise it is judged as a single-domain sequence. In DomBpred, the result is a direct output if the input sequence is predicted as a single-domain sequence. Otherwise, the full-length multi-domain sequence needs to be further decomposed.

**Figure 2.**
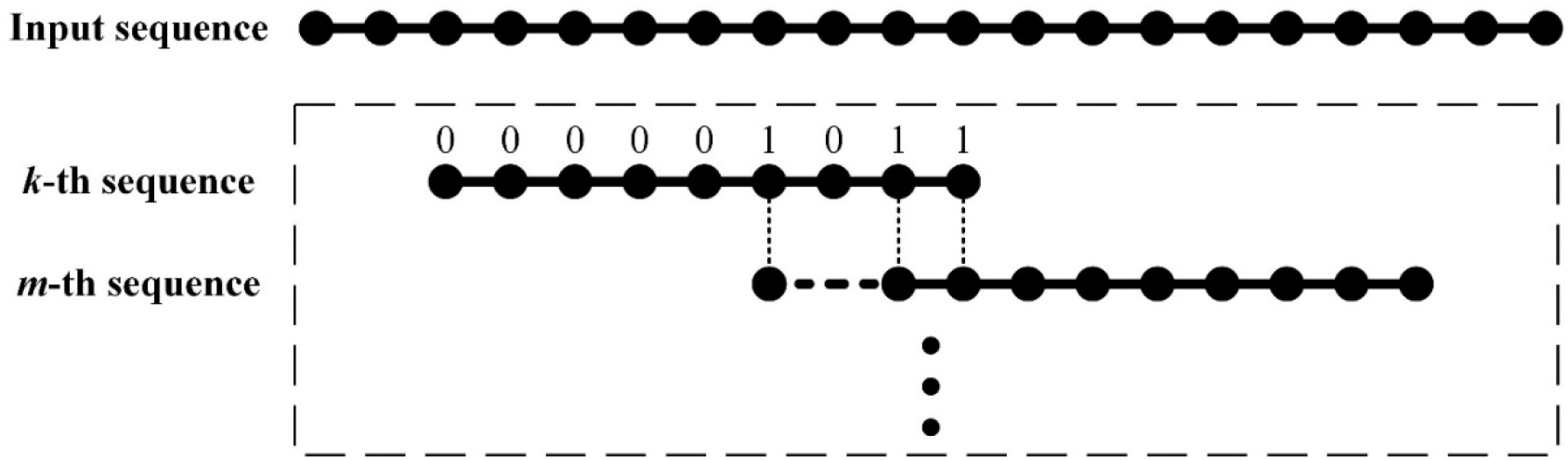
Illustration of the matched residue pair and *Mat*(*m, k*_*i*_) calculation process. When a matched residue pair is formed between *i*-th residue of the *k*-th sequence and a certain residue of the *m*-th sequence, then *Mat*(*m, k*_*i*_) = 1, otherwise *Mat*(*m, k*_*i*_) = 0. When the sum of the scores of all residues of the *k*-th sequence exceeds half of the length of the *k*-th sequence, 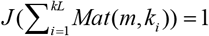; otherwise, 0.

### 2.2 Domain-residue level clustering for potential cut points

In general, a domain is a compact and separate region in a protein, and the possibility of a domain boundary being located in the compact region is low. Inspired by the Ising model, a domain-residue level clustering method is designed to cluster spatially close residues based on the distance map to form a compact region. According to the predicted distance map of the target sequence, DomBpred clusters the residues with close distances into one cluster or class, and the unclassified residues and the residues at the edge of the cluster are tuned based on the predicted secondary structure to form a set of potential cut points.

#### 2.2.1 Residue tag and update

Each residue in the protein chain is assigned a numeric tag. Herein, for a target sequence, the numeric tags are set along the protein residues increment assignment. This setup can reflect an expected characteristic, that is, the residues adjacent in sequence tend to belong to the same domain. If a residue is surrounded by neighbours with a higher tag on average, then its tag increases; otherwise, it decreases. It is defined as an update cycle when the tag of each residue in the target sequence is updated once. At each update cycle, the new tag of residue *i* is determined by its surrounding residues. The tag update formula is calculated as follows:

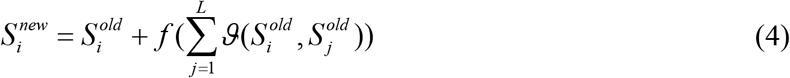

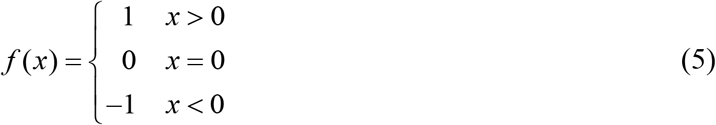

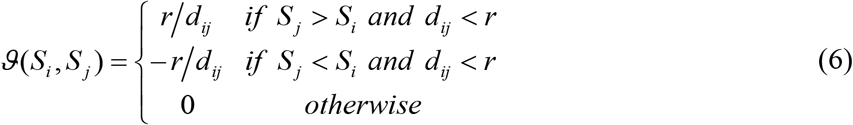

where *r* is the radius cut-off of the surrounding residues, and *d*_*ij*_ is the distance between the C_β_ atoms of residues *i* and *j* in the distance map, where the distance map is generated by trRosetta [32]. *S*_*i*_ and *S*_*j*_ represent tags of residues *i* and *j*, respectively. 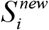 represents the tag of residue *i* after the update, and 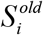 represents the tag of residue *ii* before the update.

#### 2.2.2 Dynamic neighborhood radius cut-off

The cut-off of the neighbourhood radius *r* is essential for domain-residue level clustering algorithm, and it affects the quality of potential cut points. When *r* is small, the cut points may be generated in compact regions, and the quality of these cut points is usually not high, thereby also increasing the burden for further boundary determination. When *r* is large, some spatially close domains merge together. In this case, the quality of these cut points is generally low, which also affects the accuracy of the final domain decomposition. Therefore, we design an adaptive method to dynamically determine the cut-off of the neighbourhood radius *r* according to the surrounding residues of each residue. For a residue, *r* is determined based on the residue density, and the radius corresponding to the median of the density is selected as the value of *r*. An example for the selection of *r* is shown in Supplementary Table S1. The residue density can be calculated as follows:

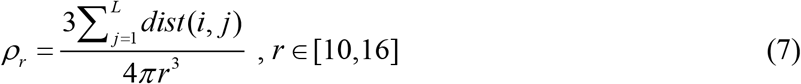

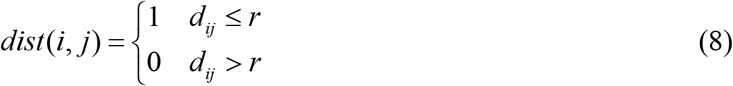

This adaptive method can ensure the quality of potential cut points, reduce the burden for further processing, and ensure that the spatially close domains are not merged together to the greatest extent.

#### 2.2.3 Iteration termination

Under the guidance of the tag update formula (4) to (6), the compact region in the sequence evolves towards the same tag. Generally, the update is stopped when the tag of the residue in the sequence does not change after an update cycle. However, the tag at the boundary of the structure domain may fluctuate, so directly determining whether to stop updating by the change in tag in one update cycle is unsuitable. Here, to terminate the update, we design an iteration termination strategy as shown in Algorithm 1. In Algorithm 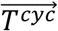 represents the tags of all residues in the input sequence after the *cyc*-th update cycle, and the cut-off parameter (*c*) is set to 10^−3^.

##### Algorithm 1 Iteration termination strategy

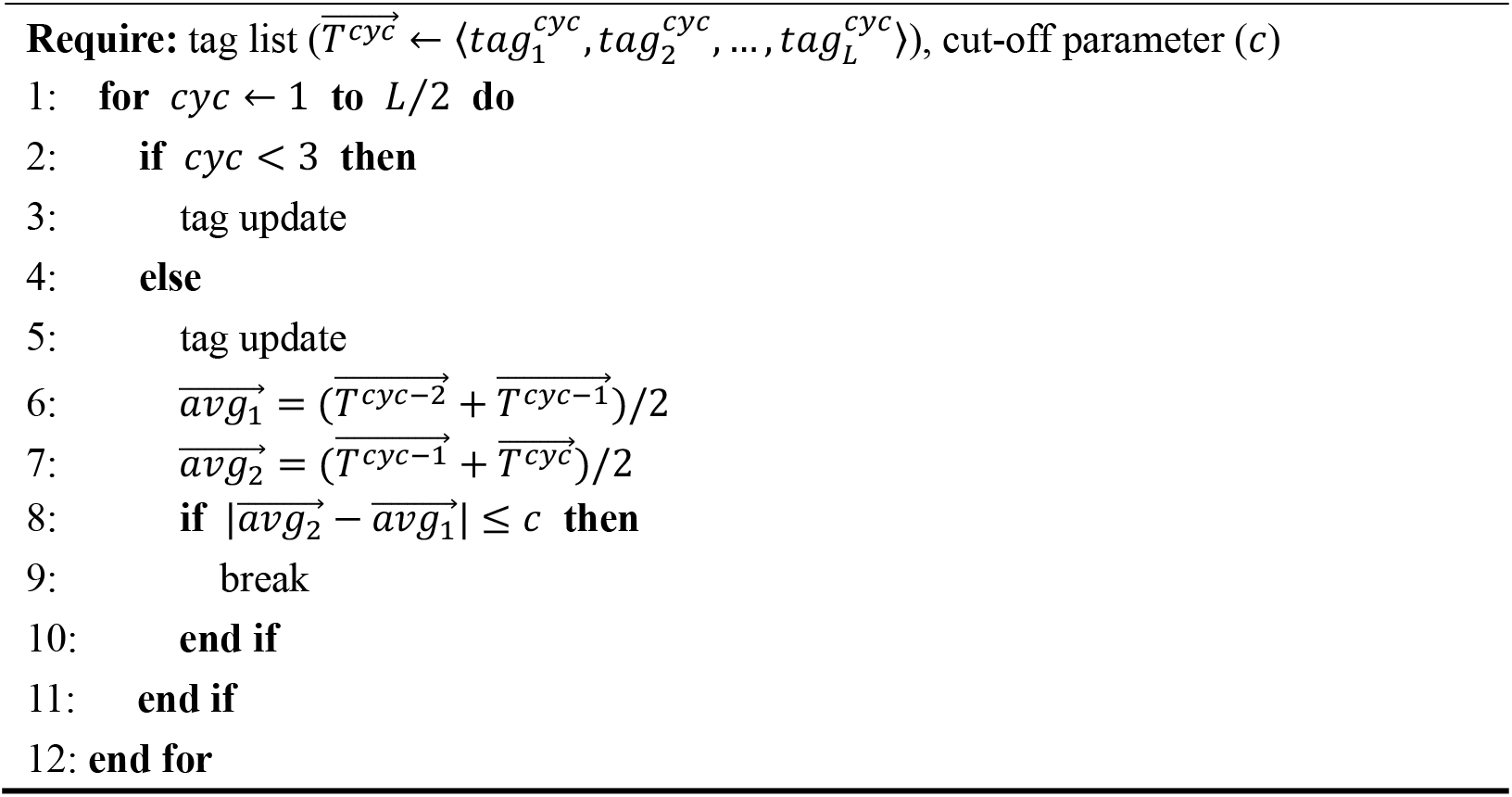

#### 2.2.4 Tag adjustment

In some proteins, the segment of one domain may be closer to other domains in space. In the DomBpred clustering process, these segments are clustered into other domains, which reduces the quality of potential cut points. Thus, we correct the segment by resetting the tags of these segments to tags of adjacent domains in the sequence. Figure 3 shows an example (1AGRA) to illustrate the process of tag adjustment. After the tag update, the protein is clustered into two regions. The residues in the dashed frame are clustered into red region because it is relatively closer to red region. However, these residues and green region belong to a same domain in the SCOPe annotation. These tags of the residues are reset to the tags of green region after the tag adjustment.

**Figure 3.**
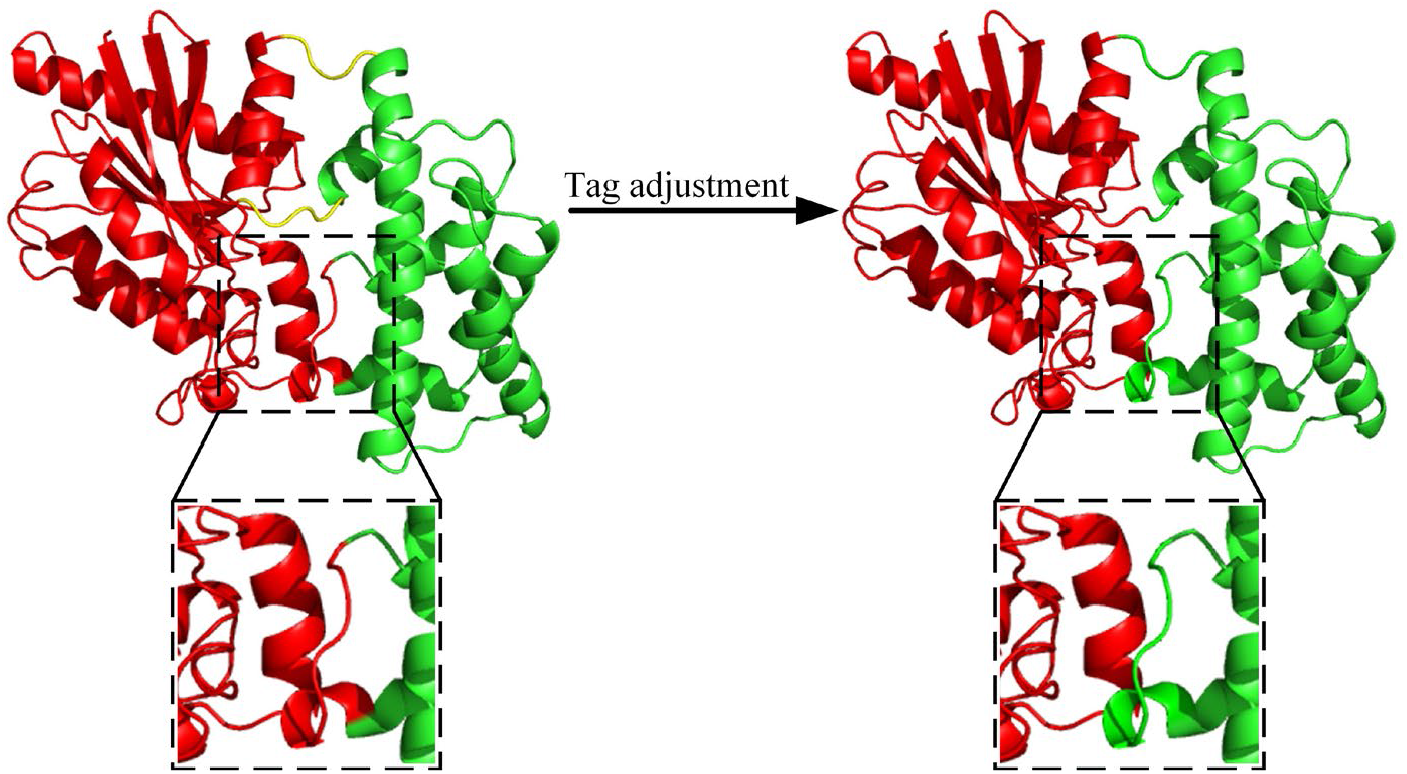
Example of tag adjustment for protein 1AGRA. Residues with the same tag are clustered into one region, such as red and green regions, whereas residues with fluctuated tags are marked in yellow. The tag of the residues in the dashed frame changed from red region to green region after the tag adjustment process.

#### 2.2.5 Potential cut point collection

As described above, the close residues on the distance map evolve towards the same tag. Therefore, adjacent residues with different tags are considered as potential cut points. The initial potential cut point set obtained by the above process needs to be further filtered. When a potential cut point is located in the *α*-helix or *β*-sheet region, the cut point is empirically inappropriate because the possibility of the domain boundary being located in the loop region is obviously higher than that in the secondary structure. In this method, when the potential cut point is located in the predicted *α*-helix or predicted *β*-sheet region, the cut point is moved to the residue of the loop region, which is the closest to the original potential cut point on the sequence. Finally, the final set of potential cut points is obtained, where the secondary structure is predicted by PSIPRED [33].

### 2.3 Domain boundary determination

The final potential cut points set may contain incorrect cut points. Thus, the potential cut points set need to be further filtered.

#### 2.3.1 Domain boundary scoring function

In general, protein domains are defined as structurally compact and separate regions of the macromolecules. A compact region can be described as a region with a large residue density, whereas a separate region can be described as a region with a small interface with other regions. Here, a domain boundary score function (*DS*) is designed to recursively evaluate whether the potential cut points are the boundary of the domain, defined as follows:

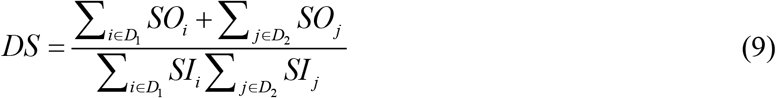

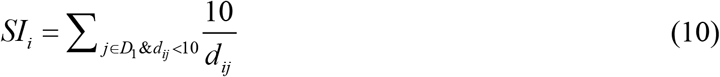

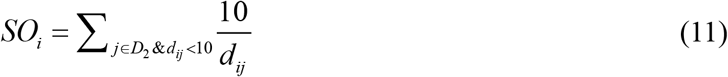

where *SI*_*i*_ is the internal compactness score of residue *i* of the first region *D*_1_ relative to *D*_1_, and *SO*_*i*_ is the external compactness score of residue *i* relative to the second region *D*_2_, where *D*_1_ and *D*_2_ are the two regions that are decomposed at the cut point.

The result of a decomposition may comprise only continuous regions or a mixture of discontinuous regions and continuous regions. Although the same *DS* score threshold can be used to measure the quality of different decomposition results, using different *DS* score thresholds may be more appropriate. Here, the two cut-off parameters *CDS*_*C*_ (cut-off for continuous domains) and *CDS*_*d*_ (cut-off for discontinuous domains) are used to judge the decomposition result comprising only continuous regions or mixed regions. These two thresholds are trained on a training set, and two optimal thresholds are obtained based on the balance of the normalised domain overlap (NDO) scores and the domain boundary distance (DBD) scores of the decomposition results.

#### 2.3.2 Domain boundary determination

The process of the domain decomposition method is shown in Figure 1 (D). Firstly, the input sequence is decomposed in a continuous domain way, that is, the input sequence is divided into two continuous domains at a potential cut point, and the *DS* of the potential cut point is calculated and denoted as *DS*_*c*_. Secondly, the input sequence is decomposed in a discontinuous domain way, that is, the input sequence is divided into a continuous and a discontinuous domain based on two potential cut points, and the *DS* of the potential cut points is denoted as *DS*_*d*_. If the *DS*_*c*_ and *DS*_*d*_ scores of the input sequence are greater than *CDS*_*C*_ and *CDS*_*d*_, respectively, that is, neither of the two ways can obtain a compact and separate domain, the input sequence is a domain and cannot be further decomposed again. If the *DS*_*c*_ or *DS*_*d*_ score of the input sequence is less than *CDS*_*C*_ or *CDS*_*d*_, one of the two decomposition ways can obtain compact and independent domains. If the *DS*_*c*_ and *DS*_*d*_ scores of the input sequence are less than *CDS*_*C*_ and *CDS*_*d*_ respectively, the input sequence is decomposed in the way with the largest difference to the corresponding cut-off. The above decomposition situation is summarised in Table 1. For the cut points in the potential cut points set, DomBpred recursively decomposes the target sequence through the above two decomposition methods until the sequence cannot be further decomposed.

**Table 1.** Summary of various decomposition situations.

| Condition | Decomposition way |
| --- | --- |
| $DS_c < CDS_c$ & $DS_d \geq CDS_d$ | decomposition in continuous domain way |
| $DS_c \geq CDS_c$ & $DS_d < CDS_d$ | decomposition in discontinuous domain way |
| $DS_c < CDS_c$ & $DS_d < CDS_d$ | decomposition with the way of largest difference |
| $DS_c \geq CDS_c$ & $DS_d \geq CDS_d$ | cannot be decomposed |

## 3 Result

In this section, we test the performance of DomBpred on the test set dataset, where its performance was compared with the threading-based method ThreaDomEx [24], and three machine learning-based methods, including FUpred [3], ConDo [27], and DoBo [26]. Notably, FUpred uses a predicted contact map to predict the protein domain boundary, and ConDo utilises contact map information as an input feature for neural network training. Meanwhile, DoBo predicts domain boundary utilising sequence and sequence profile information as the input features.

### 3.1 Datasets and assessment metrics

To fairly compare the performance of our method with other methods on the same level, the data set of FUpred [3] (including the training and test sets) is used as the data set of DomBpred, which includes 3400 single-domain and 1698 multi-domain proteins. The multi-domain proteins include 1494 continuous and 204 discontinuous domain proteins. The test set contains 849 multi-domain proteins (716 continuous and 133 discontinuous) and 1700 single-domain proteins, and the training set contains 849 multi-domain proteins (778 continuous and 71 discontinuous) and 1700 single-domain proteins.

The NDO [34] and the DBD [35] scores, used to assess domain splitting in the CASP experiments, are utilised to assess the domain boundary prediction. The NDO score calculates the overlap between the predicted and true domain regions, whereas the DBD score is defined as the distance of the predicted domain boundary from the true domain boundary, where all linker regions of the domains are considered as the true boundaries.

### 3.2 Classification of single- and multi-domain proteins

Single-domain and multi-domain proteins are two classifications of proteins. Here, we compare the domain classification capabilities of DomBpred and other four state-of-the-art methods, and the comparison results are shown in Table 2. The results of the four comparison methods originate from a published paper [3]. In the test set of 849 multi-domain proteins and 1700 single-domain proteins, the accuracy of domain classification of DomBpred is 0.945, which is 3.9% higher than that of the second-best method (FUpred). Amongst all five predictors, DomBpred generates the highest MCC (0.882), followed by FUpred (0.799), ThreaDomEx (0.759), ConDo (0.671) and DoBo (0.371). In terms of multi-domain recall, DoBo has a higher recall than DomBpred. The 23 multi-domain proteins in the test set are predicted by DoBo to be single-domain proteins, resulting in high multi-domain recall and high single-domain precision. However, the multi-domain precision and single-domain recall of DoBo are low, indicating that DoBo tends to predict the proteins as multi-domain proteins.

**Table 2.** Single- and multi-domain classification results on 2549 test proteins.

| Methods | Multi-domain |  |  | Single-domain |  |  | All |  |
| --- | --- | --- | --- | --- | --- | --- | --- | --- |
|  | Pre | Rec | F<sub>1</sub> | Pre | Rec | F<sub>1</sub> | ACC | MCC |
| DomBpred | **0.881** | 0.966 | **0.921** | **0.982** | **0.935** | **0.958** | **0.945** | **0.882** |
| FUpred | 0.860 | 0.873 | 0.867 | 0.936 | 0.929 | 0.933 | 0.910 | 0.799 |
| ThreaDomEx | 0.767 | 0.933 | 0.842 | 0.962 | 0.858 | 0.907 | 0.883 | 0.759 |
| ConDo | 0.803 | 0.751 | 0.776 | 0.880 | 0.908 | 0.894 | 0.856 | 0.671 |
| DoBo | 0.436 | **0.973** | 0.602 | 0.965 | 0.371 | 0.536 | 0.571 | 0.371 |

*Note:* Pre(multi)/Pre(single) represent the precision of multi-domain/single-domain classification and Rec(multi)/Rec(single) symbolize the recall of multi-domain/single-domain classification. F<sub>1</sub> is the harmonic average of Pre and Rec, which takes into account both. The ACC and MCC are the accuracy of protein classification and the Matthew's correlation coefficient, respectively.

In domain classification performance, DomBpred shows a better performance in overall. This is attributed to the construction of SDSL and the design of effective sequence metric. Here, DomBpred classifies multi-domain and single-domain protein by using domain knowledge in SDSL because domains are the fundamental units of protein.

### 3.3 Prediction of structural domain boundary

To examine the ability of various methods to predict the domain boundary, we present in Table 3 a summary of the NDO and DBD scores for DomBpred compared with the other four methods [3, 24, 26, 27], and the detail information for the methods on each test protein is shown in Supplementary Table S2. The NDO and DBD scores for DomBpred are higher than those of the other four methods with *P*-values < 0.05 as determined by paired one-sided Student’s *t*-tests. For the 849 multi-domain proteins in the test set, the NDO of DomBpred is 4.2%, 8.4%, 11.1% and 45.1% higher than that of FUpred, ThreaDomEx, ConDo and DoBo, respectively. Under the different NDO cut-off, the comparison of the number of the results with the meeting NDO cut-off is shown in Figure 4, and the details are shown in Supplementary Table S3. In the 849 proteins, DomBpred has 648 proteins with NDO > 0.7, whereas the best comparison method FUpred has 579 proteins with NDO > 0.7. The detailed NDO score comparison between DomBpred and the other four methods is shown in Supplementary Figure S1. The DBD of DomBpred is 5.0%, 11.0%, 39.1% and 155.1% higher than that of FUpred, ThreaDomEx, ConDo and DoBo, respectively. Under the different DBD cut-off, the comparison of the number of the results with the meeting DBD cut-off is shown in Figure 5, and the details are shown in Supplementary Table S4. The detailed DBD score comparison between DomBpred and the other four methods is shown in Supplementary Figure S2. Additionally, ConDo, ThreaDomEx, and FUpred are roughly equivalent in terms of NDO scores, but ThreaDomEx and FUpred are at least 20% higher than ConDo for DBD scores, indicating that the domain boundary predicted by ConDo is much worse than those of ThreaDomEx and FUpred. For DoBo, it performs the worst in domain boundary prediction because DoBo tends to recognize most proteins as multi-domain proteins.

**Table 3.**
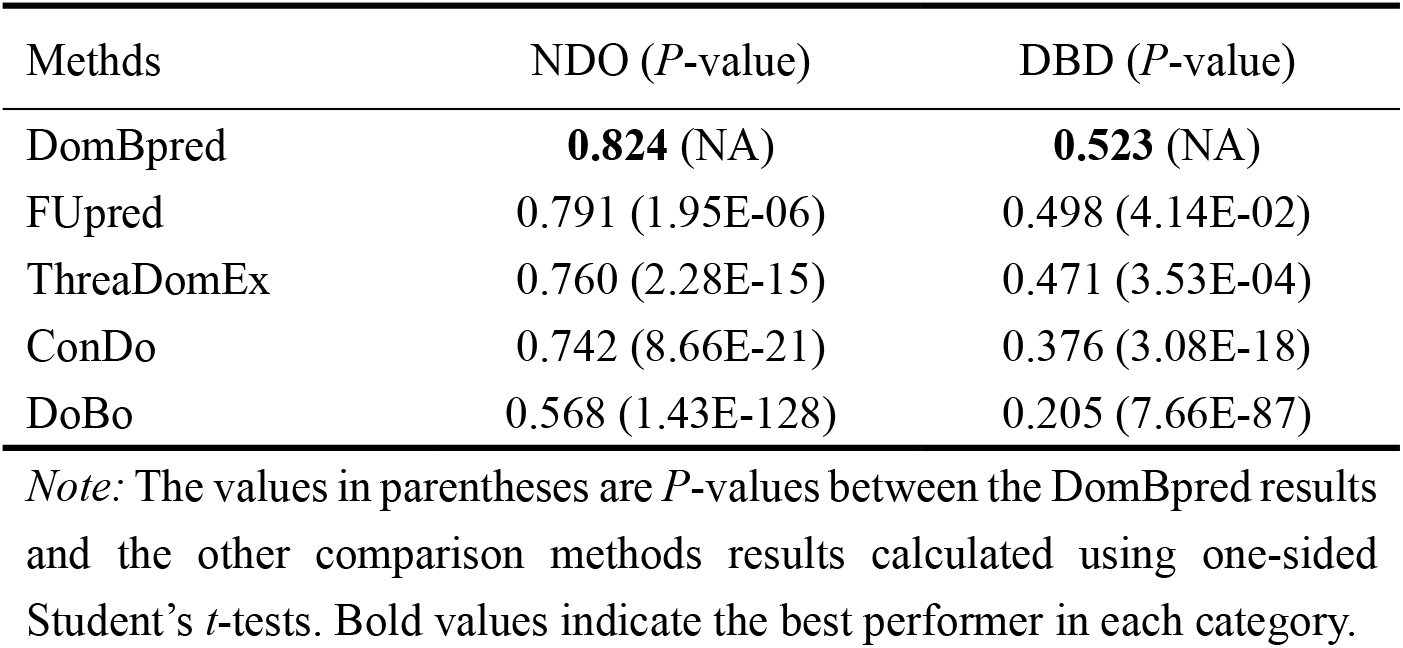
Summary of prediction results of multi-domain proteins in the test set.

**Figure 4.**
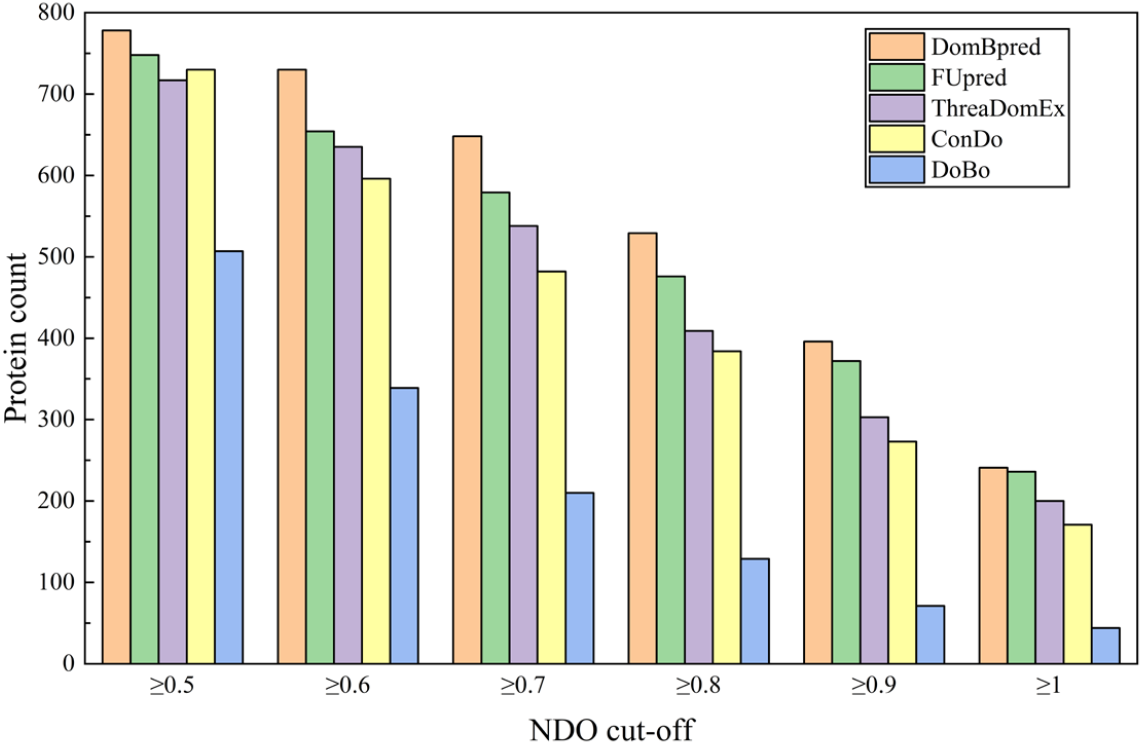
Comparison of protein quantity under different NDO cut-off. The y-axis represents the number of proteins whose NDO scores predicted by different predictors on 849 proteins meet the threshold.

**Figure 5.**
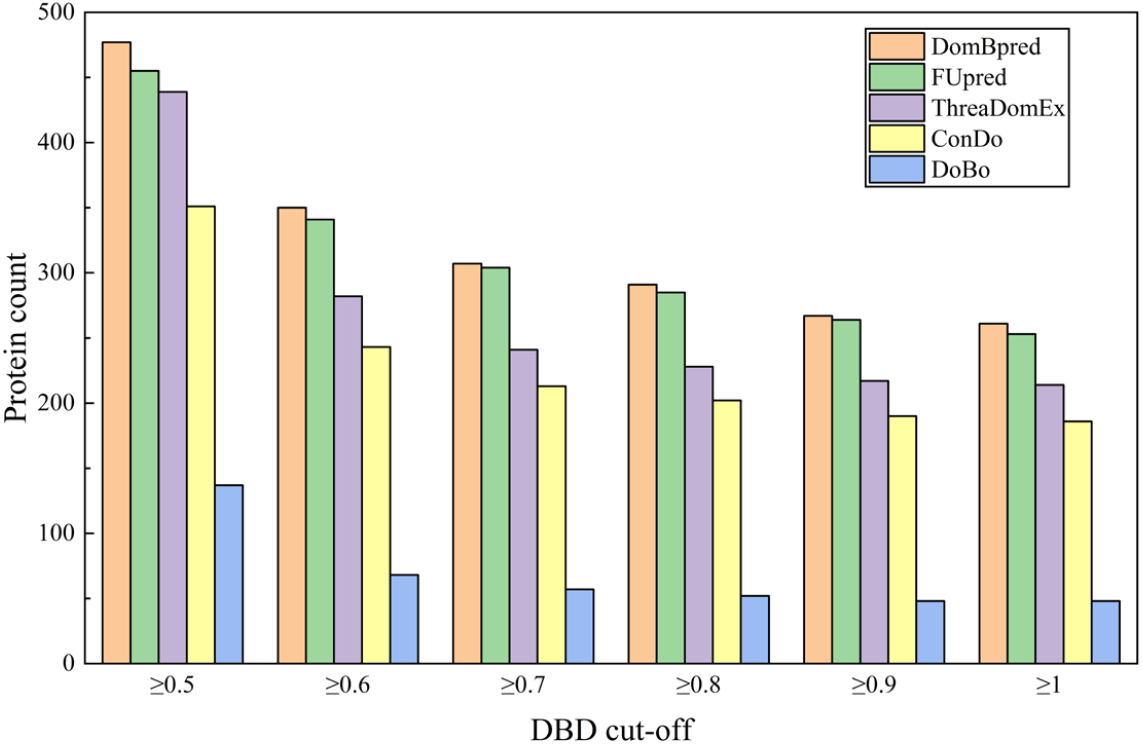
Comparison of protein quantity under different DBD cut-off. The y-axis represents the number of proteins whose DBD scores predicted by different predictors on 849 proteins meet the threshold.

These methods represent a representative set of methods on homology and machine learning based approaches, where the results demonstrate the advantage and efficiency of the DomBpred on accurately predicating the protein domain boundaries.

### 3.4 Prediction of discontinuous domain proteins

Discontinuous domains, which comprise segments from separated sequence regions, are more difficult to predict than continuous domains. Here, the prediction performance of DomBpred and the other four methods are tested on 133 discontinuous multidomain proteins in the test data set, and the results are shown in Supplementary Table S5, and the detail information is shown in Supplementary Table S6.

As shown in Table S5, amongst the 133 discontinuous multi-domain proteins, DomBpred and FUpred detect 76.7% and 70.7% of the targets containing discontinuous domains, respectively. The overall accuracy of DomBpred is lower than that of FUpred, but the NDO and DBD scores of DomBpred and that of FUpred are not significant, with *P*-value > 0.05. In terms of domain number prediction, the results of DomBpred, FUpred and ThreaDomEx are 3.05, 2.95 and 3.45, respectively. Compared with the actual number of 2.81, FUpred and DomBpred have no tendency to over-predict. The performance of DomBpred and FUpred are comparable in the discontinuous domain. The detailed NDO and DBD scores comparison between DomBpred and the other four methods is shown in Figure 6 and 7.

**Figure 6.**
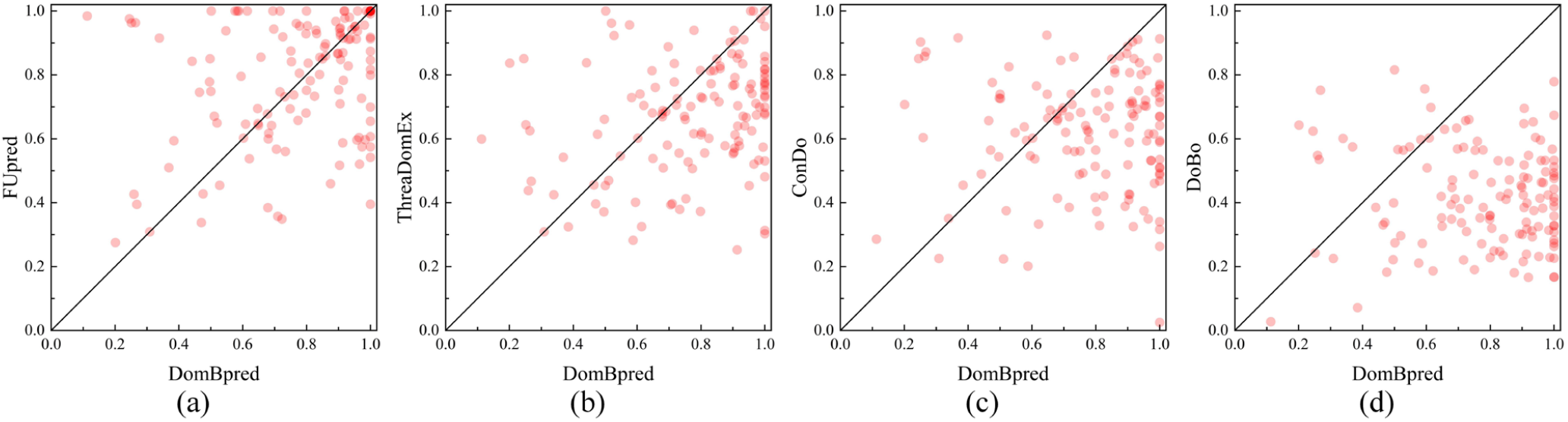
Comparison between NDO scores of DomBpred and that of other four methods on 133 discontinuous multi-domain proteins. (a) shows the comparison between DomBpred and FUpred, where the x coordinate of the red circle represents the NDO score obtained by DomBpred, and the y coordinate represents the NDO score obtained by FUpred. Similarly, (b), (c) and (d) represent the comparison between the NDO scores of DomBpred and those of ThreaDomEx, ConDo and DoBo, respectively.

**Figure 7.**
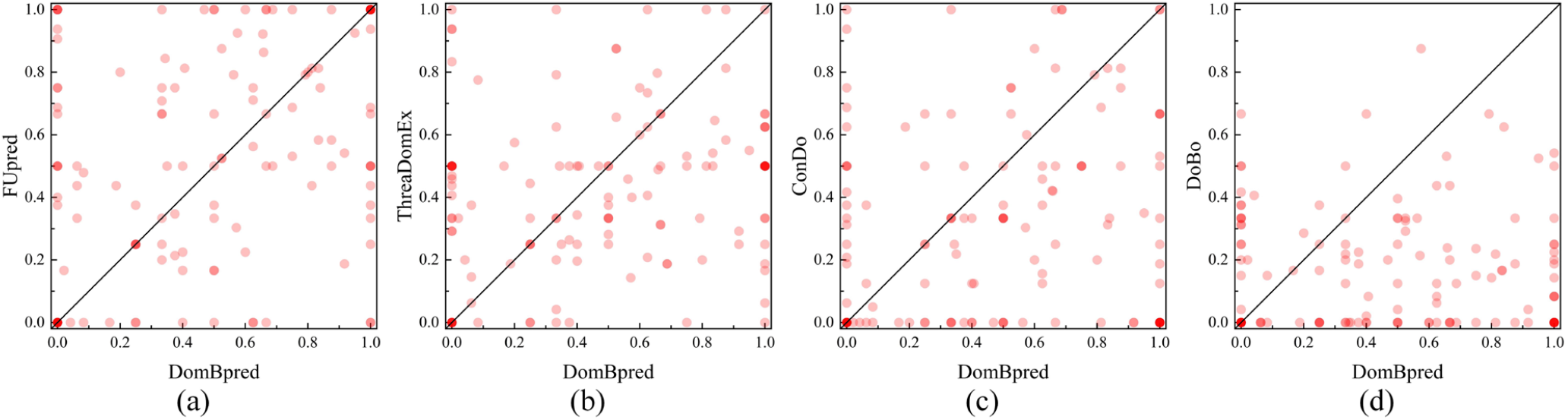
Comparison between DBD scores of DomBpred and that of other four methods on 133 discontinuous multi-domain proteins.

As mentioned above, unlike FUpred detecting domain boundary by traversing each residue of the input sequence, DomBpred detects the boundary of the domain by clustering close residues in the distance map to obtain potential cut points. Therefore, DomBpred may lose some key cut points, leading to a slight decrease in the prediction accuracy of the discontinuous domain, but reducing the computational cost of the detection domain boundary. Moreover, ConDo and DoBo cannot detect any proteins containing discontinuous domains.

### 3.5 Dynamic radius

We compare the performance of DomBpred under different neighbourhood cut-off radius on the test set, and the results are shown in Table 4. The dynamic radius achieves better results in all cases. It is because DomBpred can cluster relatively reasonable compact regions under the dynamic radius strategy, which can detect better potential cut points. Here, the protein (PDB ID: 1L5JA) with a length of 862 residues is taken as an example to illustrate the relationship between clustering of close residues and the detection of potential cut points under different radius.

**Table 4.** Summary of the prediction results of multi-domain proteins in the test set under different radius.

| Different radius (Å) | NDO | DBD |
| --- | --- | --- |
| Dynamic ([10,16]) | **0.8335** | **0.5510** |
| 10 | 0.8262 | 0.5390 |
| 11 | 0.8286 | 0.5346 |
| 12 | 0.8292 | 0.5421 |
| 13 | 0.8286 | 0.5442 |
| 14 | 0.8318 | 0.5451 |
| 15 | 0.8302 | 0.5488 |
| 16 | 0.8231 | 0.5384 |

The different decomposition results of 1L5JA are shown in Figure 8. The number of potential cut points for fixed radius 10, 11, 12, 13, 14, 15, 16 and dynamic radius are 30, 21, 22, 24, 13, 3, 7 and 11, respectively. DomBpred with a radius of 10 detects 30 potential cut points, resulting that the three-domain protein is divided into six-domain proteins. By contrast, DomBpred with a radius of 16 detects 7 potential cut points, and there is no cut point for the second domain, resulting that the three-domain protein has the wrong domain boundary. DomBpred with dynamic radius detects 11 potential cut points. Finally, the protein is divided into four-domain proteins, in which the third domain determined by SCOPe is divided into two domains by DomBpred. However, it is also appropriate to divide into two domains from the perspective of compactness. Although it does not completely agree with the annotations, it also achieves the highest scores compared with fixed radius. This shows that the dynamic radius can also balance the ability to detect potential cut points whilst clustering compact areas as much as possible, rather than tending to one side.

**Figure 8.**
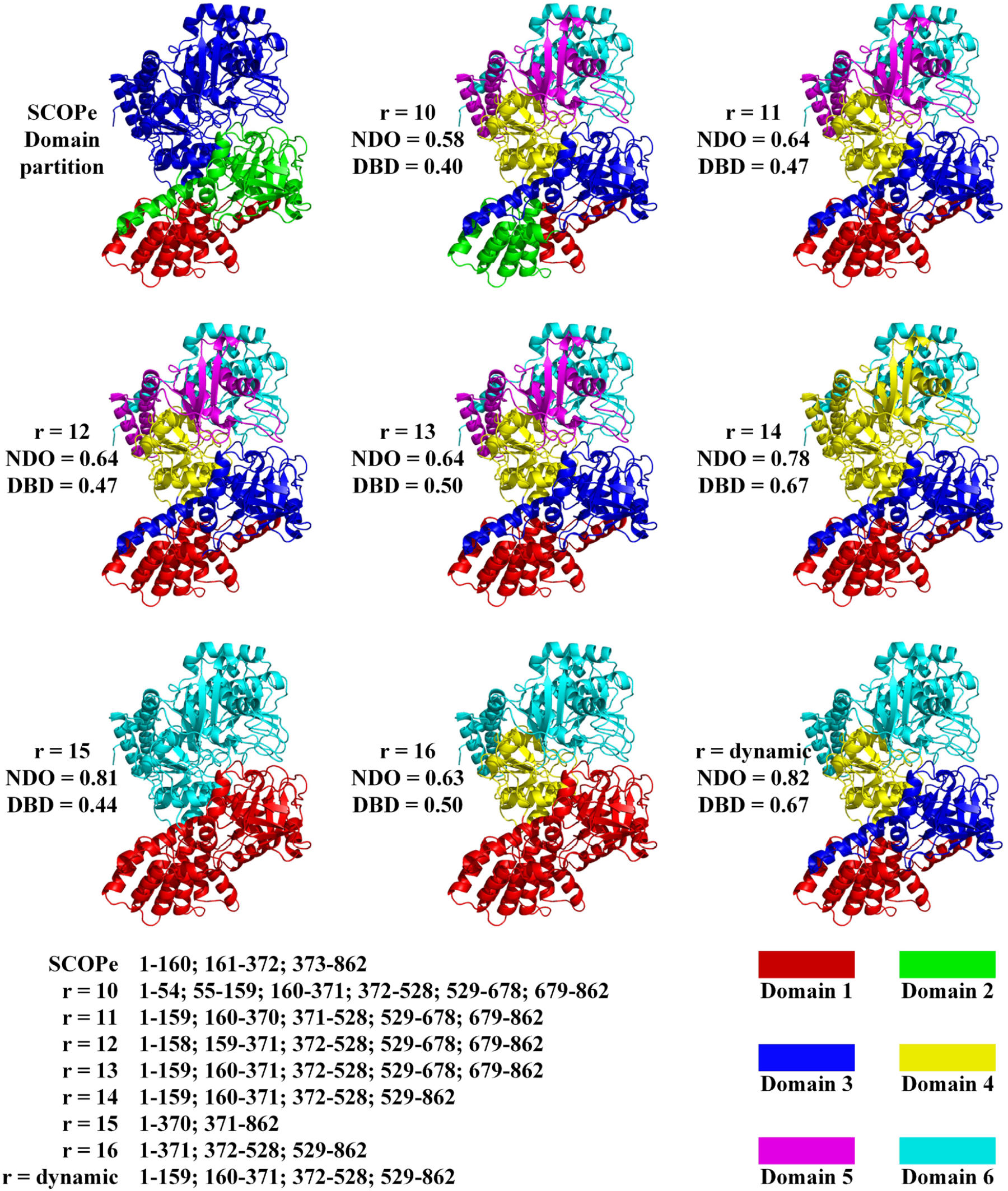
Case study of domain prediction of E. coli aconitase protein (PDB ID: 1L5JA) under different radius. Under different radius, the NDO and DBD scores of the domain prediction results are listed on the left side of the 3D structure diagram, and the specific decomposition results are listed at the bottom of the 3D diagram.

### 3.6 What went right and wrong

On one hand, the homology-based methods can have a high accuracy of predictions when close templates are identified. Accordingly, DomBpred is inspired by ideas from this homology-based method, and then constructs a SDSL to initially identify the classification of input sequence. As a result, 1564 single-domain sequences of 1700 are correctly classified. The 25 sequences of the remaining 136 single-domain sequences are further correctly classified in the following process of DomBpred. Finally, DomBpred achieves precision with 98.2% in 1700 single-domain sequences, as shown in Table 2. On the other hand, the accuracy of the sequence-based method using machine learning is improved with the development of the machine learning. Compared to contact information, the distance information may better contribute to predict the domain boundary. Therefore, DomBpred uses the distance map information to detect the boundary of the structure domain. DomBpred detects the domain boundary by clustering close residues in the distance map to obtain potential cut points. This may make DomBpred filter out some pseudo cut point residues. Therefore, DomBpred better performs on 849 multi-domain proteins.

In the 133 discontinuous proteins amongst 849 proteins, the NDO and DBD of DomBpred are lower than those of FUpred, but no significant difference exists between them. The results may be due to the fact that some potential cut points at key positions are omitted because DomBpred obtains potential cut points by clustering close residues in the distance map, leading to decreased prediction accuracy of the discontinuous protein. However, FUpred traverses every residue, which may explain why the accuracy of DomBpred is not as good as that of FUpred in discontinuous proteins.

## 4. Conclusion

We developed a sequence-based protein domain predictor, named DomBpred, using inter-residue distance and domain-residue level clustering inspired by Ising model to predict the protein domain boundary. In DomBpred, we construct a comprehensive domain sequence database based on SCOPe and CATH databases, and an effective sequence metric is proposed to detect the classification of input sequence. At the same time, a clustering method inspired by Ising model is proposed to cluster the close residues in the distance map to form a cluster, in which the residues are considered to be located in a compact region. For the unclassified residues in the clustering results and the residues at the edge of the compact regions, these residues are further evaluated using the designed domain boundary scoring function to identify the domain boundary. Furthermore, the dynamic radius strategy is used to determine the range of close residues, which can avoid the irrationality caused by the fixed radius to some extent.

DomBpred is compared with FUpred, ThreaDomEx, ConDo and DoBo in 2549 test set proteins. The experimental results on the given test set show that the overall performace of DomBpred is better than those of existing approaches. In 2549 test proteins, DomBpred generated correct single-and multi-domain classifications with a Matthew’s correlation coefficient of 0.882. In 849 multi-domain proteins, the DBD and NDO scores of DomBpred are 0.523 and 0.824, respectively, which are 5.0% and 4.2% higher than those of the best comparison method. Notably, the proposed method depends to some extent on the accuracy of distance map. With the advancement of machine learning in distance map prediction, a higher-accuracy distance map may further improve the prediction accuracy of DomBpred.

**Key points**

- We design a clustering method for target sequence, which uses inter-residue distance and domain-residue level clustering algorithm inspired by Ising model to cluster spatially close residues into compact regions.
- We construct a single-domain sequence library (SDSL) and propose an effective sequence metric to identify single-domain and multi-domain proteins.
- Based on domain independence and compactness, a domain boundary score function is designed to select the boundary points.
- Results of the comparison of test set proteins suggest that our proposed domain boundary prediction method outperforms four other state-of-the-art full-version methods.

## Supplementary Data

Supplementary data are available online at BIB.

## Data and code availability

All data needed to evaluate the conclusions are present in the paper and the Supplementary Materials. The additional data and code related to this paper can be downloaded from https://github.com/iobio-zjut/DomBpred.

