## Supplementary Material for "DomBpred: protein domain boundary predictor using inter-residue distance and domain-residue level clustering"

### S1. Dynamic neighborhood radius cut-off

**Table S1.** Example of residue neighborhood cut-off radius selection.

| $r$ | 10 | 11 | 12 | 13 | 14 | **15** | 16 |
|---|---|---|---|---|---|---|---|
| $\rho(r)$ | 0.191 | 0.179 | 0.152 | 0.131 | 0.122 | **0.141** | 0.105 |
| Ranking | 1 | 2 | 3 | 5 | 6 | **4** | 7 |

The $\rho(r)$ row in the Table S1 represents the surrounding residue density of a residue with $r$ as the neighborhood radius, for example, the surrounding residue density of a residue with 10 as the neighborhood radius is 0.191.

The Ranking row in the Table S1 represents the ranking of the surrounding residue density of a residue under different neighborhood radius. For example, the surrounding residue density of a residue with 15 as the neighborhood radius ranks fourth.

In DomBpred, the neighborhood radius of a residue is the radius corresponding to the median value of the surrounding residue density. In Table S1, DomBpred will select 15 as the neighborhood radius of this residue, because the surrounding residue density corresponding to the radius 15 is in the middle of all the densities.

### S2. Other tables and figures

**Table S3.** Comparison of protein quantity under different NDO cut-off on the 849 proteins in the test set.

| Methods | ≥0.5 | ≥0.6 | ≥0.7 | ≥0.8 | ≥0.9 | ≥1.0 |
|---|---|---|---|---|---|---|
| DomBpred | 778 | 730 | 648 | 529 | 396 | 241 |
| FUpred | 748 | 654 | 579 | 476 | 372 | 236 |
| ThreaDomEx | 717 | 635 | 538 | 409 | 303 | 200 |
| ConDo | 730 | 596 | 482 | 384 | 273 | 171 |
| DoBo | 507 | 339 | 210 | 129 | 71 | 44 |

**Table S4.** Comparison of protein quantity under different DBD cut-off on the 849 proteins in the test set.

| Methods | ≥0.5 | ≥0.6 | ≥0.7 | ≥0.8 | ≥0.9 | ≥1.0 |
|---|---|---|---|---|---|---|
| DomBpred | 477 | 350 | 307 | 291 | 267 | 261 |
| FUpred | 455 | 341 | 304 | 285 | 264 | 253 |
| ThreaDomEx | 439 | 282 | 241 | 228 | 217 | 214 |
| ConDo | 351 | 243 | 213 | 202 | 190 | 186 |
| DoBo | 137 | 68 | 57 | 52 | 48 | 48 |

**Table S5.** Summary of prediction results of 133 discontinuous multi-domain proteins in the test set.

| Methods | NDO (*P*-value) | DBD (*P*-value) | Recall |
|---|---|---|---|
| DomBpred | 0.772 (NA) | 0.475 (NA) | **102** |
| FUpred | **0.788** (2.20E-01) | **0.521** (9.01E-02) | 94 |
| ThreaDomEx | 0.672 (2.27E-06) | 0.421 (5.88E-02) | 52 |
| ConDo | 0.620 (3.94E-09) | 0.312 (1.62E-05) | 0 |
| DoBo | 0.417 (5.99E-30) | 0.171 (1.30E-14) | 0 |

*Note:* The values in parentheses are *P*-values between the DomBpred results and the other control methods results calculated using one-sided Student's *t*-tests. Bold values indicate the best performer in each category.

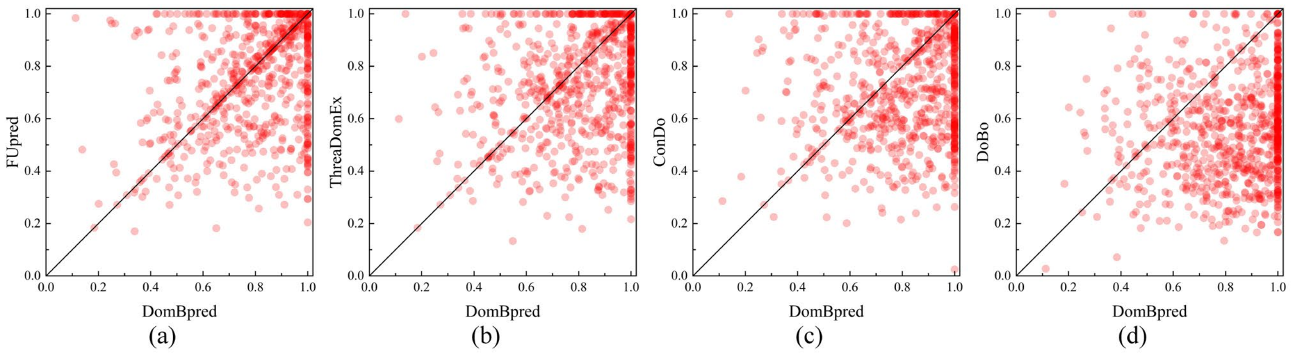


**Figure S1.** Comparison between NDO scores of DomBpred and that of other four methods on 849 multi-domain proteins. Figure S1 (a) shows the comparison between DomBpred and FUpred, where the x coordinate of the red circle represents the NDO score obtained through DomBpred, and the y coordinate represents the NDO score obtained through FUpred. Similarly, Figure S1 (b), (c) and (d) represent the comparison between the NDO scores of DomBpred and that of ThreaDomEx, ConDo and DoBo, respectively.

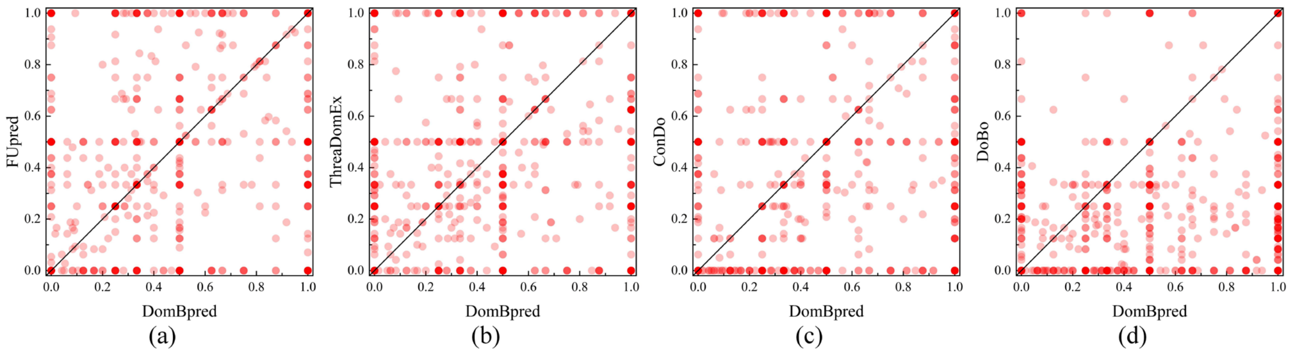


**Figure S2.** Comparison between DBD scores of DomBpred and that of other four methods on 849 multi-domain proteins. Figure S2 (a) shows the comparison between DomBpred and FUpred, where the x coordinate of the red circle represents the DBD score obtained through DomBpred, and the y coordinate represents the DBD score obtained through FUpred. Similarly, Figure S2 (b), (c) and (d) represent the comparison between the DBD scores of DomBpred and that of ThreaDomEx, ConDo and DoBo, respectively.
